# Repurposing Live Attenuated Trivalent MMR Vaccine as Cost-effective Cancer Immunotherapy

**DOI:** 10.1101/2022.02.25.481685

**Authors:** Yuguo Zhang, Musa Gabere, Mika Taylor, Camila C. Simoes, Chelsae Dumbauld, Oumar Barro, Jean Christopher Chamcheu, Steven R. Post, Thomas J. Kelly, Mitesh J. Borad, Martin J. Cannon, Alexei Basnakian, Bolni M. Nagalo

## Abstract

Despite its rising promise, cancer immunotherapy remains out of reach for many patients because of the extensive cost of manufacturing immunotherapy products. In this study, we show that intratumoral injections of the trivalent measles, mumps, and rubella (MMR) live attenuated viral vaccine (LAVs) modulates a potent cytotoxic T-cell antitumor immune response, resulting in tumor growth inhibition and improved survival in syngeneic mouse models of hepatocellular carcinoma and colorectal cancer. Using an integrated transcriptomic and proteomic approach, we demonstrated that mechanistically, MMR exerts its antitumor activity by priming innate and adaptive antitumor immune responses, leading to immunologically coordinated cancer cells death. Our findings highlight a promising potential for LAVs, such as MMR, to be repurposed as cost-effective cancer immunotherapy.

## INTRODUCTION

Hepatocellular carcinoma (HCC) is one of the most frequently diagnosed cancers in the world’s most disadvantaged areas.^1-3^ HCC is responsible for 90% of all primary liver cancers,^4^ annually claiming approximately 800,000 lives worldwide.^4-7^ With all available therapies, the 5-year survival rate is less than 15% for advanced unresectable HCC.^8^ While hepatitis B virus and hepatitis C virus are the most prominent risk factors for HCC,^9^ in recent decades, HCC prevalence has tripled in the US due to the increased incidence of metabolic risk factors for liver diseases, chiefly nonalcoholic steatohepatitis, nonalcoholic fatty liver disease, obesity, and type II diabetes.^10-13^ Therapeutic approaches for early-stage HCC include hepatic resection, transplantation, and thermoablation for small lesions.^14^ Unfortunately, less than 30% of HCC patients are eligible for these therapies due to poor baseline liver function and high tumor burden.^15,16^

In the past few years, immunotherapy has gained momentum in cancer treatment by enabling improvements to patient survival.^14,17-21^ Oncolytic viral therapy is an emerging new class of cancer immunotherapy that involves selectively infecting and killing tumor cells.^22-25^ Oncolytic viruses (OVs) induce versatile and multimodal immune responses involving innate and adaptive immunity.^26,27^ This unique ability, shared among viruses and bacteria, has been exploited to develop vaccines and anticancer agents.^28^ Several OVs have already exhibited promising potential for treating human cancers.^29-32^ In spite of the evidence of therapeutic benefit,^33,34^ seamless clinical use of OVs in cancer patients also faces numerous challenges. These limitations comprise the varying efficacy of OVs in human cancers,^4^ and more importantly, the high cost of their development and manufacturing, which makes them far out of reach for patients with lower socioeconomic status.^35^ Because of these reasons, developing economically sustainable analogous approaches to current high-cost OVs is an area of high interest in oncology.

Studies have suggested that one exciting approach to expand the clinical benefit of OVs to many patients is to repurpose live attenuated viral vaccines (LAVs) for cancer immunotherapy.^36,37^ But these early studies have been limited by their focus on LAVs made up of a single viral vector. ^36-41^ As others have shown, multiple mechanisms of tumor resistance, including changes in antitumor immune response pathways, enable cancer cells to become resistant to the antitumor effect of OVs.^42-44^ Therefore, we postulate that targeting cancer cells with different lineages of OVs can elicit distinct, specific, and robust immune responses with the potential to alleviate these therapeutic resistances.

Therefore, we capitalize on the immunostimulatory activity of the trivalent measles, mumps, and rubella live vaccine (MMR) to delineate a cost-effective cancer immunotherapy. MMR is attractive as a virotherapy primarily for its proven safety record but has other useful features as well. MMR has been shown to stimulate a potent and long-lasting protective immunity against measles, mumps, and rubella in humans.^45-50^ And multiple studies have also demonstrated that vaccine lineages of measles and mumps viruses are effective against human tumors in cell culture, animal models, and clinical trials.^38,40,41,51,52^ Furthermore, MMR is readily available, low-cost, and accessible worldwide, and its use requires no regulatory approval from the US Food and Drug Administration (FDA).

Here, we report that intratumoral (IT) injections of MMR elicit a robust antitumor cytotoxic T-cell immune response, leading to tumor regression and extended survival in animal models of HCC and colorectal cancer (CRC). We selected these two models because the liver is one of the dominant metastatic sites for colorectal cancer cells.^53,54^ Measles, mumps, and rubella have been shown to not productively replicate in mouse cancer cells.^55-58^ And OVs promote antitumor immunity against infected and uninfected cancer cells.^43,59-62^ Thus, we employed an integrated transcriptomic and proteomic approach to conclusively elucidate the mechanisms of the antitumor immune response of MMR in murine tumors. Overall, these findings demonstrate that MMR modulates a potent antitumor immune response, warranting its further evaluation as potential cost-effective cancer immunotherapy.

## RESULTS

### IT injections of MMR induce subcutaneous tumor growth inhibition and extend survival in mouse tumor models

Based on the abundant clinical evidence of the immunomodulatory properties of MMR,^45-50^ we hypothesize that its IT administrations can elicit an antitumor immune response. To investigate this, we subcutaneously implanted mouse HCC (Hepa 1-6) or CRC (MC38) cells into the right flanks of immune-competent mice. At days 0, 7, and 14 mice were administered 50 µL IT doses of phosphate-buffered saline (PBS; PBS control groups) or a low-dose MMR (1 × 10^2^ TCID_50_ for each virus; MMR groups) (Supplementary Fig. 1). We found that extended dosing of MMR resulted in significant tumor inhibition and extended survival compared to that in the control group (PBS) in Hepa 1-6 (tumor, *p=*0.001; survival, *p=*0.04) and MC38 (tumor, *p=*0.03; survival, *p=*0.02) models (Figs. 1a and 1b).

**Figure 1.**
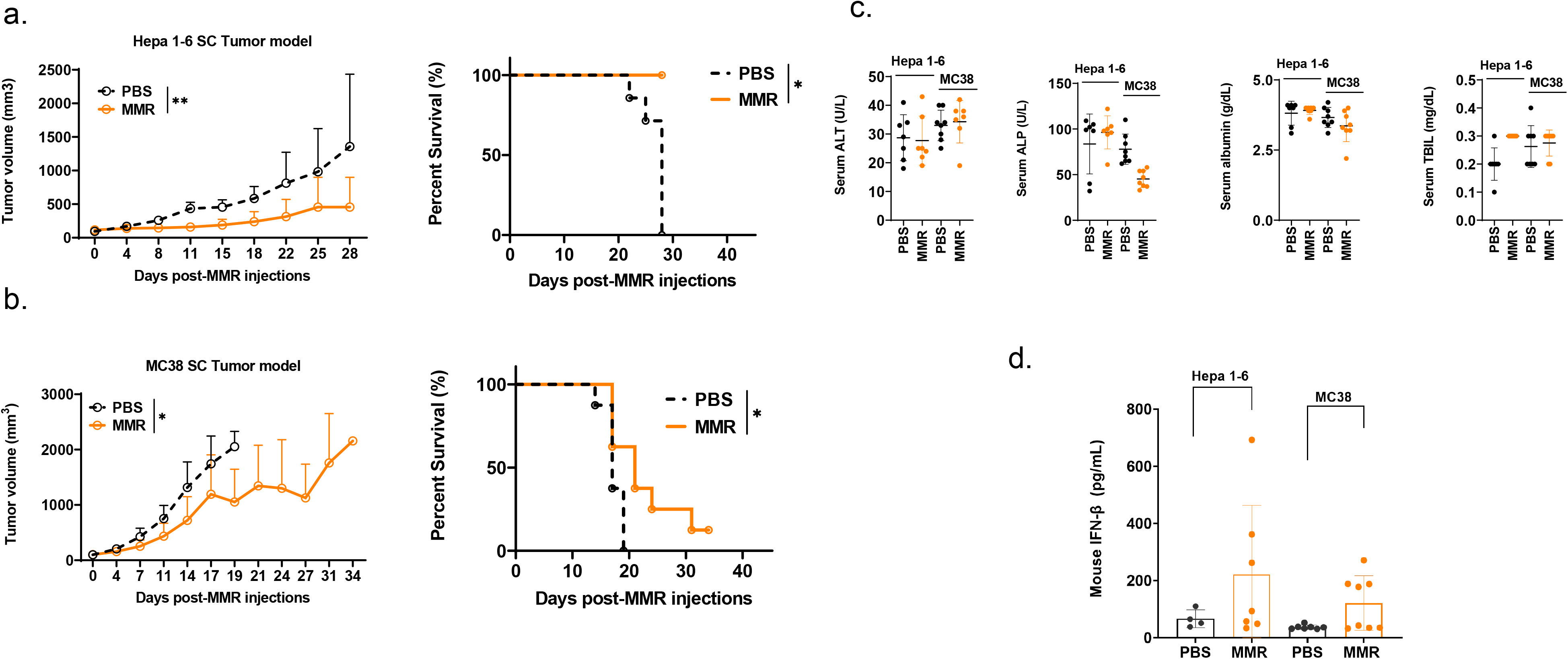
Intratumoral injections of MMR induce tumor inhibition and extend survival in two murine tumor models. **(a) and (b). Tumor growth and survival curves**. Hepa 1-6 (n=7/group) or MC38 (n=8/group) were allowed to grow on the right flanks of immune-competent mice. Once tumors reached 80-120 mm^3^, mice were intratumorally administered low doses (1 × 10^2^ TCID_50_) of the measles, mumps, and rubella (MMR) live vaccine or phosphate-buffered saline (PBS; controls). Tumor volumes (mm^3^) were measured twice weekly using a digital caliper and plotted as a function of time. Panel (**a)** and **(b)** show changes in tumor volumes over time and Kaplan-Meier curves for survival for Hepa 1-6 and MC38 tumor models, respectively. The day when we injected the first MMR or PBS into the mice is defined as day 0. The statistical significance of differences in the survival curves between the groups was evaluated using the log-rank (Mantel-Cox) test. **(c) Mouse Serum biochemical analysis**. Graphs showing changes in liver function tests, including concentration of enzymes such as alkaline phosphates (ALP), alanine aminotransferase (ALT), and total bilirubin (TBIL) between the PBS and MMR-treated groups. **(d)** *Levels of mouse type I interferon β (IFN-β) in serum*. Level of antiviral cytokine (IFN-β) was measured in serum from mice treated with measles, mumps, and rubella (MMR) live vaccine or phosphate-buffered saline (PBS; controls).

In comparison, the IT dose of MMR used in this study is significantly (10^3^-10^4^-Fold) lower than that of oncolytic measles and mumps (i.e., 10^7^-10^8^ TCID_50_) in mouse tumor models and clinical trials. ^38,40,41,51,52^ This extra safety feature of MMR makes it very attractive because the administration of a high dose of viral vectors has been shown to induce severe liver and brain toxicity. ^63^ Furthermore, terminal serum chemistry analysis showed no difference in the levels of markers for liver toxicity (i.e., alanine aminotransferase, alkaline phosphatase, total bilirubin) or nephrotoxicity (i.e., urea nitrogen and creatine) in the two groups (PBS and MMR), indicating that IT injections of MMR are well tolerated in mice (Fig. 1c and Supplementary Fig. 2). MMR induced expression of mouse type I interferon, indicating activation of antiviral immune mechanisms (Fig. 1d), and no change in body weight was observed (Supplementary Fig. 3).^64^ Moreover, the antitumor effect of MMR was not significantly impacted by seroconversion to measles and mumps (Supplementary Figs. 4a and 4b). Interestingly, we did not detect antibodies to the rubella virus in our samples (Supplementary Fig. 4c). Human vaccine studies have reported a broad spectrum of differences in serum antibodies levels to rubella among vaccinated individuals, including waning or low antibodies responses.^65^ Therefore, the negative results could also be due to the wanning or low levels of anti-rubella antibodies.

### Measles, mumps, and rubella viruses infect and lyse Vero cells, but do not replicate in murine tumors

To determine if MMR forms plaques on producer cells, we infected a monolayer of Vero cells with a low dose of MMR (1 × 10^2^ TCID_50_ for each virus) in a 6-well plate for four days. After the incubation period, MMR-infected cells were washed with PBS and fixed with 10% paraformaldehyde for 10 minutes, and representative plaques were visualized by crystal violet staining (Supplementary Fig. 5a). To assess whether MMR-induced antitumor activity is due to viral replication and oncolysis, we amplified viral genes using specific primers to genes coding for measles nucleoprotein, mumps matrix protein, rubella envelope protein and murine β-actin in Hepa 1-6 and MC38 tumors. However, our amplification did not detect viral genes in Hepa 1-6 and MC38 tumors, but murine β-actin was successfully amplified (Supplementary Fig. 5b). These results suggest that MMR can efficiently infect and lyse producer cells and trigger a virus-induced oncolysis-independent antitumor activity and improved survival in murine models of HCC and CRC without the clinical pathology effect observed with other oncolytic viruses (i.e., hepatotoxicity and nephrotoxicity).^66,67^

### IT administrations of MMR increased the frequency of tumor-infiltration immune cells

To determine the various aspects of the immune response associated with IT administrations of MMR in vivo, at the end of the study, Hepa 1-6 tumors were surgically removed and dissociated into single-cell suspension (see materials and methods).^68^ A multicolor flow cytometry assay was used to identify and assess the frequency of tumor-infiltrating (TILs) immune cells (e.g., F4/80 [macrophages], CD8 [T cells], CD4 [helper T cells], CD11+ [dendritic cells], NK [NK cells]). In comparison to IT injection of PBS, IT injection of MMR was associated with increased tumor infiltration of immune cells, particularly cytotoxic (CD8+) T cells (Fig. 2a). However, only the frequency of CD8+ GranzymeB+ TILs population (*p*=0.02) was significantly upregulated in tumors treated with MMR compared to PBS (Fig. 2b.). By contrast mice that received IT injections of MMR had decreased numbers of F4/80-(Fig. 2c; *p*=0.03) and M1 macrophages (Fig. 2d: *p*=0.02), but not in the frequency of CD4+ (CD+ Ki67, CD4+ PD-1, CD4+ IFNg+) (Fig. 5). M1 macrophages have been associated with resistance to pathogens and inhibition of carcinogenesis, but the M2 phenotype exerts immunosuppressive effects.^69,70^ Although not statistically significant, there was an evident decrease in the frequency of total macrophages, including M2 phenotype and NK cells in the MMR group compared to PBS controls (Supplementary Fig. 6). Interferon-gamma (IFNg+) is a potent antiviral cytokine that is essential for cytotoxic T cell-mediated elimination of measles, mumps, and rubella viruses and the establishment of antiviral immunity. ^71-73^ In this study, we found no difference in the subsets of CD8+ IFNg+ and CD4+ IFNg+ between the MMR-treated mice and PBS-treated mice, suggesting an absence of viral replication or clearance of viral particles. CD8+ and CD4+ T lymphocytes are essential effectors of adaptive antiviral immune response.^74^ Previous studies showed that depletion of CD8+ T cells in primates exposed to wild-type measles is associated with severe disease (extensive rash, higher viral loads, and persistent viremia).^75,76^ Moreover, it has long been reported that children with a defect in T-lymphocyte activity often develop fatal diseases.^77^ Since both mumps and rubella vaccination elicit potent humoral and cellular immune responses.^72,78-80^ In the absence of viral replication^55-58^, our findings highlight the important role of IT infiltration of CD8+ T cells and decrease in the frequency of macropahges in MMR-induced antitumor activity likely via activation of class I major histocompatibility (MHC)-restricted CD8+ T cells.

**Figure 2.**
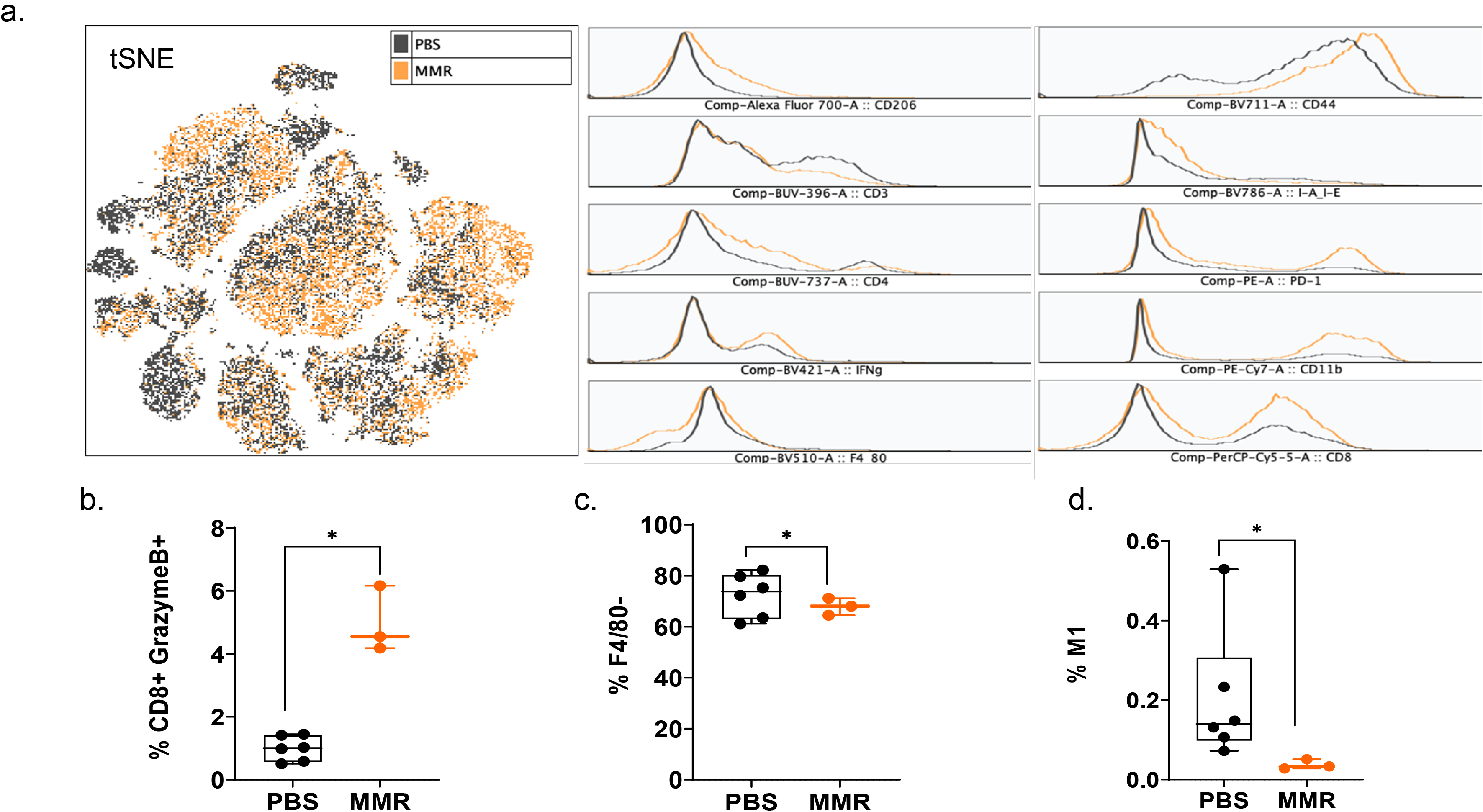
Analysis of tumor-infiltrating immune cells following intratumoral injection of the MMR vaccine. (**a**) T-distributed stochastic neighbor embedding (tSNE) analysis showed a dense intratumoral population of murine immune cells in Hepa 1-6 tumors treated with measles, mumps, and rubella (MMR) live vaccine compared to controls treated with phosphate-buffered saline (PBS). Graphs showing increased frequencies of tumor-infiltrating CD8+ granzyme B+ reactive CTLs **(b)**, F4/80-cells **(c)**, and M1 macrophages **(d)** in Hepa 1-6 tumors. The Bartlett test was used to test homogeneity of variance and normality. If the *p* value of the Bartlett test was no less than *p*=0.05, ANOVA and a two-sample *t* test were used to compare group means. If the *p* value of the Bartlett test was less than 0.05, the Kruskal-Wallis and Wilcoxon rank-sum tests were used to compare group means. The figures demonstrate the potential significant difference of the gated subsets in the mCD45+ population, determined by the *p* value.

### MMR activates innate and adaptive antitumor immune response pathways

To validate the biological relevance of the observed MMR-mediated T-cell antitumor immunity, we analyzed the transcriptome of Hepa 1-6 tumors injected with PBS or MMR. Analysis of mRNA expression levels of these two groups using the limma-voom method^81^ identified a storm of 4,331 upregulated and 3,126 downregulated genes (Fig. 3a); however, the expression of 15,329 genes was relatively unaffected. The volcano plot shows the gene expression difference distribution (Fig. 3b). Based on this result, we performed Gene Set Enrichment Analysis (GSEA) on the differentially expressed genes (DEG) between the PBS and MMR groups. We applied two functions (gseGO and gseKEGG) to identify the enriched terms in Gene Ontology (GO) and the Kyoto Encyclopedia of Genes and Genomes (KEGG) with a false discovery rate (FDR) value < 0.05. The top biological functions and canonical pathways that were up-or down-regulated by MMR were activation of the immune response, adaptive immune response, immune response-activating cells surface receptor signaling pathway, immunoglobulin production, and production of molecular mediator of immune response (Fig. 3c). Furthermore, the KEGG analysis indicated that the differentially expressed genes were mainly involved in immunoglobulin complex and circulating immunoglobulin complex, which are crucial in the regulation of antigen-mediated immune response (Supplementary Fig. 7). These results demonstrate that mechanistically, IT administrations of MMR trigger local antigen presentation, leading to activation and recruitment of cytotoxic effectors of innate and adaptive immunity, which are the hallmarks of a durable antitumor immune response.^82-87^

**Figure 3.**
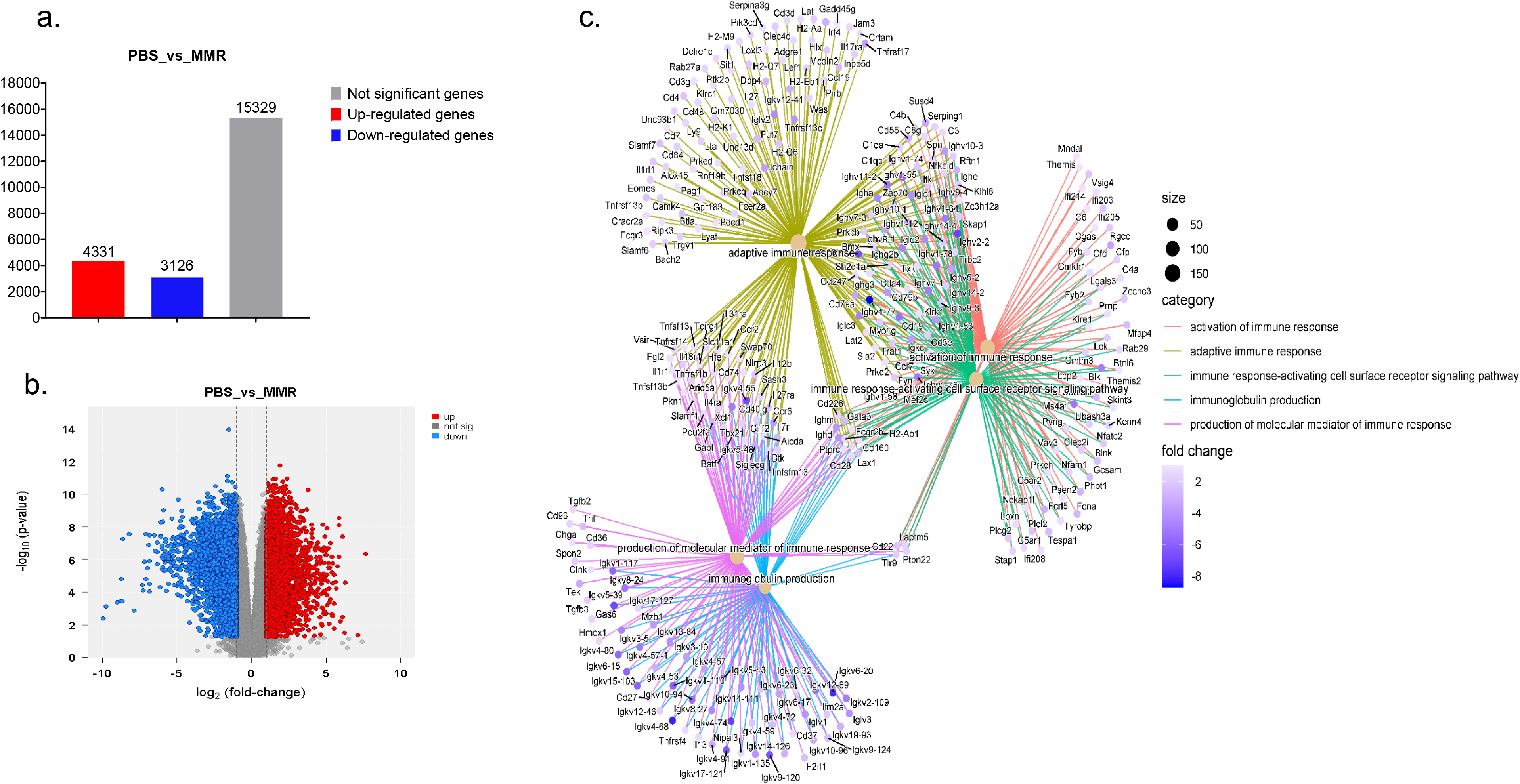
Transcriptomic analysis of tumor treated with the MMR vaccine. To determine the mechanism of antitumor activity of the measles, mumps, and rubella (MMR) live vaccine in vivo, we measured mRNA expression using transcriptomic in Hepa 1-6 mice tumors. Differentially expressed genes were determined using the limma-voom method. A fold-change |*log*FC| ≥ 1 and false discovery rate (FDR) of 0.055 were used as a cutoff. The *log*FC was computed using the difference of the mean of *log2*(MMR) and mean of *log-2*(PBS), that is, mean of *log2*(MMR) - mean of *log2*(PBS). **(a)** Graphs showing the total number of genes upregulated (red bar), (down-regulated (blue), and unchanged (gray) between the PBS vs. MMR groups. **(b)** Shows a volcano plot indicating the gene expression difference distribution. **(c) Gene ontology (GO) analysis using gseGO**. Visualization of Gene Set Enrichment Analysis (GSEA) on the DEG between PBS controls and MMR groups. The significant differences are shown in some graph, where *, P-value < 0.05; **, P-value < 0.01; ***, P-value < 0.001.

### Integrated analysis of transcriptomic and proteomic approaches identified proteogenomic changes differences associated with MMR in murine HCC

We applied a correlation analysis to examine the association between mRNA and protein expression levels in the two groups (PBS and MMR) of Hepa 1-6 tumors. First, a MixOmics supervised analysis was used to select the best features from the multi-omics data to differentiate mRNAs and corresponding expressed proteins between the PBS and MMR groups. We identified the top 30 enriched features differentially expressed in the PBS group compared to the MMR group (Fig. 4). The top normalized mRNA and protein intensities useful in discriminating MMR from the PBS cohort are listed in an integrated dataset heatmap (Supplementary Fig. 8).

**Figure 4.**
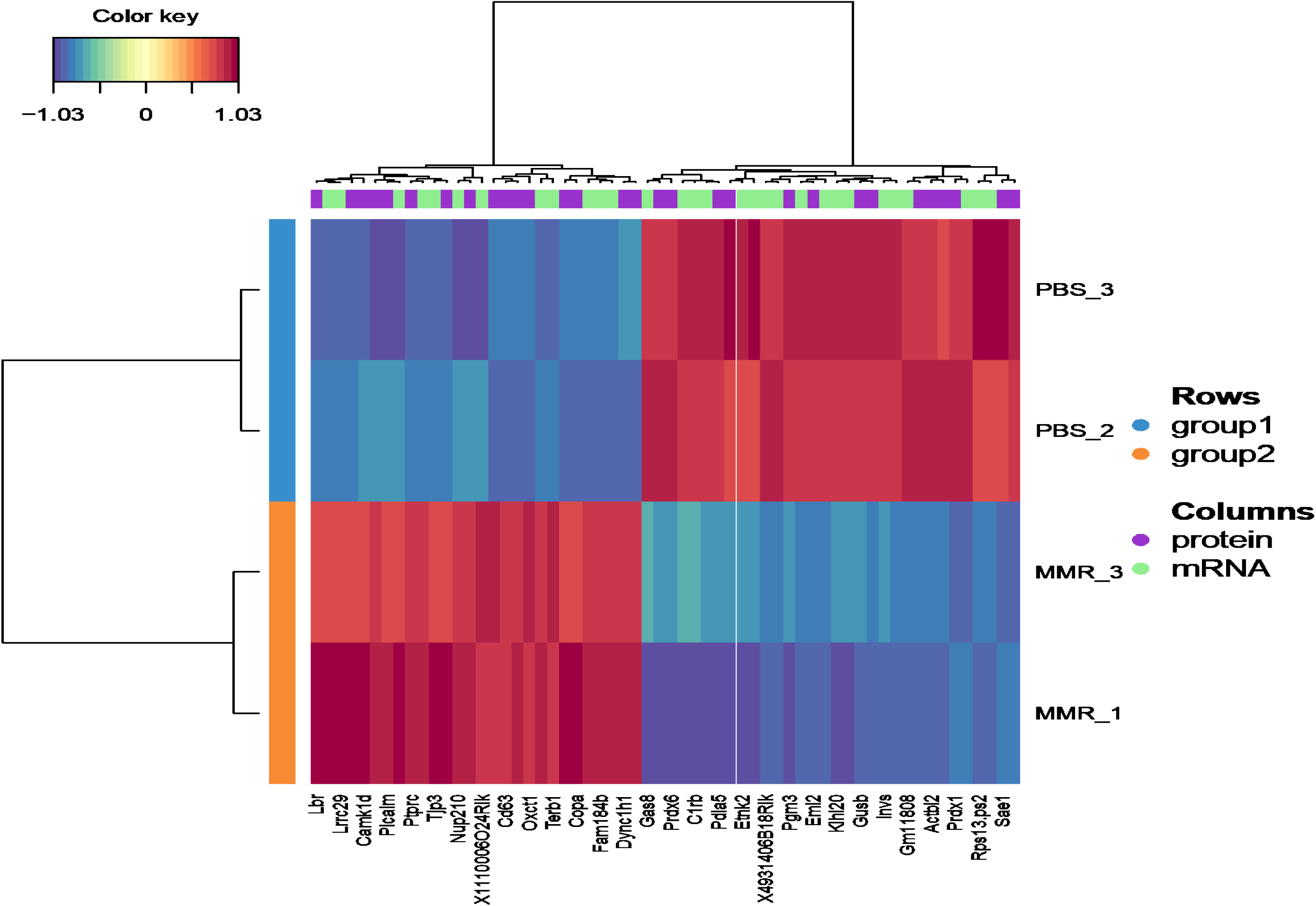
Integrated Analysis of protein intensities and RNA expression in Hepa 1-6 tumors treated with MMR. A MixOmics supervised analysis was used to display the top genes/proteins contributing to the difference in integrated datasets in a heatmap.

Out of the differentially expressed mRNA intensities that correlate with their protein counterparts, we identified the dynein cytoplasmic 1 heavy chain, an essential member of the intracellular transport of DNA damage proteins family with a controversial role in cancer. Indeed, in humans, while its downregulation has been linked to poor prognosis and low survival in glioblastoma^88^, its upregulation has been shown to delay tumor proliferation in gastrointestinal tumor cells.^89^ Similarly, the coatomer subunit α (COPA) gene encodes the human homolog of the a-subunit coatomer protein complex involved in intracellular protein transport. Loss of COPA has also been associated with the high proliferation and invasiveness of HCC cells.^90^ Others features, such as CD63, were identified, which overexpresses negatively regulated tumor invasiveness, including HCC.^91^ Furthermore, several of these genes and their protein products, such as PTPRC, NUP210, OXCT1, ETNK2, are known to be upregulated in tumors cells, including HCC.^92-95^ Others are involved in cellular senescence, apoptosis, angiogenesis, inflammation, and wound healing (LBR, EML2, SAE1).^96-99^ Some of these genes (i.e., CAMK1D, TJP3, GAS8, and PDIA5) are associated with epithelial-mesenchymal transition (EMT), immune cells infiltration, and immune evasion.^100,101^

Analysis of top KEGG pathways also indicated an upregulation of Leukocyte transendothelial migration, endocytosis (phagocytosis), neutrophil extracellular traps signaling (cellular death) pathways, and downregulation of pathways like viral carcinogenesis and drug metabolism, suggesting an MMR-induced activation of inflammation, innate and adaptive immune responses (Supplementary Fig. 9). A dataset of genes that contributed most to the biological process (phagocytosis and cellular death) highlighted in the KEGG pathways is displayed in Supplementary Fig. 10.^102^ Downregulation of other pathways such as hypoxia and reactive oxygen species, revealed by the MSigDB analysis in MMR-treated tumors, also indicated activation of cellular metabolic arrest and apoptosis (Supplementary Fig. 11). These results further validate our transcriptomic findings indicating that MMR can function as a cancer vaccine via recruitment and activation of effectors of innate and adaptive antitumor immune response pathways.

## Discussion

Studies have shown that defects in interferon (IFN) pathways favor OV-mediated tumor-restricted oncolysis.^103,104^ OVs can infect, replicate and lyse a wide range of mammalian tumor cells, leaving normal cells unaffected.^22-28^ In addition, virus-mediated oncolysis provides needful conditions for priming antitumor immunity by activation of tumor-specific cytotoxic T cells.^52,62,82-86,105-109^ To date, there are only three immunovirotherapies approved for use in human cancers, including the herpes simplex virus T-Vec, the modified adenovirus H101, and the picornavirus Rigvir,^110-114^ even though many other OVs have been evaluated clinically.^29-32,40,115-118^ This is because the cost-intensive nature of manufacturing and testing clinal grade OVs make them far out of reach for most patients.^35^ The trivalent MMR vaccine effectively modulates durable immunity against measles, mumps, and rubella, making it one of the most successful human vaccines to date.^45-50^ Numerous preclinical and clinical studies have demonstrated the high immunogenicity and antitumor efficacy of vaccine lineages of measles and mumps.^38-41^ Based upon extensive data on the immunomodulatory properties of MMR, we evaluate the antitumor potential of its IT administrations in two preclinical mouse models.

Here, we show that IT immunotherapy with low doses of MMR lead to prominent tumor growth inhibition and improved survival in subcutaneous syngeneic hepatocellular carcinoma (HCC) and colorectal cancer (CRC) models. HCC disproportionally affects patients from disadvantaged backgrounds globally,^5,119^ and the liver is one of the dominant metastatic sites for colorectal cancer cells.^53,54^ Additionally, HCC is often localized at advanced stages^120^, allowing administrations of high concentrations of viral vectors while mitigating off-target effect and the impact of virus-neutralizing antibodies (NAbs).^121-123^ Hence, why we chose to focus primarily on HCC and CRC in this study.

A plethora of studies have shown that the antitumor activity of OVs is predominantly due to the potent activation of tumor-specific T cells that recognize tumor antigens.^83,85,86,107,109^ In this study, we also show that MMR modulates such antineoplastic immunity via activation and IT infiltration of effector CD8+ T cells. Previous works have pointed out that pre-existing immunity to OVs is detrimental to their oncolytic activity.^51,124,125^ Moreover, several authors have shown that neutralizing antibodies to measles can hamper its systemic delivery and antitumor effect.^122,126^ Accordingly, strategies to overcome this limitation have been proposed with various degrees of success. ^109,121,122,127^ However, we found that the antitumor activity of IT doses of MMR is not significantly affected by seroconversion to measles and mumps viruses. So far, the preferred method of injecting OVs in patients with solid tumors is the IT route of administration.^128^ In addition, it has been shown that antibodies targeting measles hemaglutinin (H) protein can enhance its cellular uptake rather than inhibit viral entry.^129^ These results once again fortify the list of evidence validating that IT administration of immunovirotherapy can improve the safety of OVs via reducing the spread of viral vectors to normal tissues and the effect of NAbs on the bioavailability of viral vectors.

Currently, there is a scarcity of transcriptomic and proteomic data explaining the antitumor mechanisms of OVs. Therefore, we employed an integrated proteogenomic pipeline to identify mRNA and corresponding protein intensities differentially expressed following IT injections with MMR or control in murine HCC tumors. This approach allowed us to mechanistically elucidate that MMR potently primed multiple effectors involved in innate and adaptive immunity resulting in a coordinated cytotoxic T cell-mediated killing of tumor cells. Because measles, mumps, and rubella are exclusive human pathogens that poorly replicate in mouse tissues.^55-58^ Thus, to study measles and vesicular stomatitis virus pseudotyped with measles entry proteins, we and others have used a modified mouse model expressing the human measles Edmonton strain receptor complement regulatory protein (CD46).^56,130^ Hence, we amplified viral genes to investigate whether the observed tumor growth inhibition was also due to virus-induced oncolysis. Consistent with these previous reports, we found that none of these viruses extensively replicated in the murine tumors, strongly indicating that MMR promoted a lysis-independent activation of antitumor immunity likely via recruitment of class I MHC-restricted CD8+ T cells.^43,60-62,131^

This study shows that multiple low IT doses of MMR can yield similar, if not greater, anti-tumor activity than high doses of single OVs in mouse tumor models.^31,52,132,133^ Besides, while oncolytic measles and mumps have demonstrated safety in laboratory animals and humans, relatively high viral titers are required to achieve the desired antitumor effect in humans.^38-41,52,132^ This could be particularly challenging in pediatric, older, and pregnant patients. Because the administration of high titers of viral vectors in primates and humans can often result in severe hepatotoxicity and deficiency in motor neurons, often leading to death. ^63,67,134-136^ This work shows that potent in vivo antitumor efficacy can be obtained using low doses of measles, mumps, and rubella viruses. Since these viruses are antigenically different, thus a concomitant infection can elicit distinct immune responses, resulting in a much more powerful and additive antitumor activity than any of these viruses could have induced alone. More importantly, despite being in early to late phases of clinical development for more than a decade in the US, the FDA has not yet approved oncolytic measles in human cancers.^38,41,107,137^ This investigation demonstrates that IT treatments with MMR can be a safe, potent alternative, and cost-effective approach to overcome therapeutic resistance that single OVs (i.e., oncolytic measles) encounter in cancers.

Although our data indicate that MMR induces a robust antitumor immunity, it does not determine if MMR elicits memory T-cell responses.^44,138,139^ It did not also provide insights into the contribution of immune response to measles, mumps, and rubella specific antigens to the antitumor effect of MMR or the impact of virus-neutralizing antibodies on its systemic administration.^140,141^ The type and stage of cancer, immune mechanisms, timing, dosage, and route of administration are crucial for obtaining the desired therapeutic effect with OVs. One of the objectives of this work is to impart a detailed mechanistic understanding of the antitumor activity of IT administrations of MMR. This will enable the rational design of studies using MMR alone or combined with other therapies, such as immune checkpoint inhibitors, in early or late-stage solid tumors for possible additive or synergistic long-term responses in clinical settings.^37^

Lastly, we have shown that therapeutic strategies that repurpose low-cost trivalent LAVs to activate antitumor immunity are achievable with MMR. These findings can revolutionize treatment and improve clinical outcomes for economically disadvantaged cancer patients; but even so, to fully exploit the antitumor properties of MMR, future work should focus on understanding to what extent (potency and durability) MMR-enabled antitumor immunity can effectively target and eradicate minimal residual disease and distant metastases in experimental cancer models, then clinical studies.

## MATERIALS AND METHODS

### Cell lines

The murine hepatoma Hepa 1-6 (ATCC CRL-1830) and Vero (ATCC CCL-81) cell lines used in this study were purchased from ATCC and was cultured in Dulbecco’s modified eagle medium (DMEM) supplemented with 10% fetal bovine serum (FBS), 1% L-glutamine, and 1% penicillin/streptomycin. The murine colon adenocarcinoma cells MC-38 (Cat. # ENH204-FP; Kerafast) used in this study were obtained from Kerafast and cultured in DMEM with 10% FBS, 2 mM glutamine, 0.1 mM nonessential amino acids, 1 mM sodium pyruvate, 10 mM Hepes, 50 µg/mL gentamycin sulfate, and 1% penicillin/streptomycin. All cells were passaged in a tissue culture incubator at 37 °C and 5% CO_2_.

### Preparation of the measles, mumps, and rubella live vaccine

The measles, mumps, and rubella live virus vaccine (MMR) was purchased from the University of Arkansas for Medical Sciences (UAMS) pharmacy and contained attenuated live Edmonston measles, B level Jeryl Lynn mumps, and RA 27/3 Rubella viral strains. A single immunizing dose (single 500 µL vial) of the MMR vaccine delivers 1 × 10^3^, 1.25 × 10^3^, and 1 × 10^3^ median tissue culture infectious doses (TCID_50_) of measles, mumps, and rubella viruses. In this study, we used a 10-fold lower dose (1 × 10^2^, 1.25 × 10^2^, and 1 × 10^2^ TCID_50_ for each virus) compared to the immunizing dose. To prepare the vaccine for animal studies, lyophilized MMR vaccine powder vials were reconstituted and diluted with the provided diluents as recommended by the manufacturer (Merck).

### Visualization of MMR-induced plaque formation in Vero cells

We used a low dose of MMR (1 × 10^2^ TCID_50_) to infect adherent Vero cells (2.5 × 10^5^ cells per well) in 6-well plates. Cells were incubated at 37 °C until analysis. At 4 days post-infection, cells were fixed with 10% paraformaldehyde (PFA) and stained with 0.1% crystal violet to visualize virus-induced plaques in infected and mock-infected wells. We took pictures of representative areas in two MMR-infected wells and one mock infected well.

## ANIMAL STUDIES

Under an approuved UAMS Animal Use Protocol, we conducted the in vivo evaluations described below.

### In vivo efficacy of the MMR vaccine in a syngeneic HCC and CC models

To evaluate the in vivo therapeutic efficacy of the MMR vaccine in subcutaneous mouse HCC and CC models, 1 × 10^6^ Hepa 1-6 or MC38 cells in 100 µL of cold RPMI were injected subcutaneously into the right flanks of immunocompetent female C57BL/6 mice (n=7/group [Hepa 1-6], n=8/group [MC38]; Jackson Laboratories) using 1 mL syringes. Mice were monitored weekly for palpable tumors or any changes in appearance or behavior. When average tumors reached a treatable size (80 to 120 mm^3^), mice were randomized into the respective study groups and dosed within 24 hours of randomization. On days 0, 7, and 14, mice were administered 50 µL IT injections containing PBS (PBS control groups) or 1 × 10^2^ TCID_50_ units (a 10-fold lower dose compared to the immunizing dose used in children) of MMR (MMR groups). Tumor volume and body weight were measured twice weekly following randomization and initiation of treatment using a digital caliper and balance. Tumor volume was calculated using the following equation: (longest diameter * shortest diameter2)/2 with a digital caliper. During the first week of treatment and after each injections, mice were monitored daily for signs of recovery for up to 72 hours. Mice were euthanized when body weight loss exceeded 20%, when tumor size was larger than 2,000 mm^3^, or for adverse effects of treatment. Mice were sacrificed 28 (Hepa 1-6 cohort) or 34 (MC38 cohort) days following the first MMR dose administration, at which time tumor and blood were collected for downstream analysis.

### Analysis of tumor-infiltrating immune cells

Hepa 1-6 tumors (n= 6 for PBS, n=3 for MMR) were excised and dissociated using a mouse tumor dissociation kit (Miltenyi, CAT# 130-096-730) with a gentleMACS^(tm)^ Octo Dissociator (Miltenyi) according to the manufacturer’s protocol. CD45^+^ cells were isolated with mouse CD45 (TIL) microbeads (Miltenyi). Cells were incubated with Fixable Viability Stain 510 for 15 minutes at 4 °C followed by anti-Fc blocking reagent (Biolegend, Cat# 101320) for 10 minutes prior to surface staining. Cells were stained, followed by data acquisition with a BD LSRFortessa X-20 flow cytometer. All antibodies were used following the manufacturer’s recommendation. Fluorescence Minus One control was used for each independent experiment to establish gating. For intracellular staining of granzyme B, cells were stained using an intracellular staining kit (Miltenyi), and analysis was performed using FlowJo(tm) (TreeStar). Forward scatter and side scatter cytometry were used to exclude cell debris and doublets.

### Flow cytometry antibody analysis

The following antibodies were used for flow cytometry analysis: CD45-FITC (Cat. # 553079; BD Biosciences), CD3-BUV395 (Cat. # 563565; BD Biosciences), CD4-BUV737 (Cat. # 612761; BD Biosciences), CD8-Percp-Cy5.5 (Cat. # 45-0081-82; eBioscience), CD44-BV711 (Cat. # 103057; Biolegend), CD335-PE/Dazzle594 (Cat. #137630; Biolegend), PD-1-PE (Cat. # 551892; BD Biosciences), Ki67*-BV605 (Cat. # 652413; Biolegend), Granzyme B*-APC (Cat. # 366408; Biolegend), IFN-γ*-BV421 (Cat. # 563376; BD Biosciences), CD11b-PE-Cy7 (Cat. # 101216; Biolegend), F4/80-BV510 (Cat. # 123135;, Biolegend), CD206-AF700 (Cat. # 141734; Biolegend), I-A/I-E-BV786 (Cat. # 743875; BD Biosciences), and L/D-efluor780 (Cat. # 65-0865-18; eBioscience).

### Proteomic analysis of HCC formalin-fixed paraffin-embedded tissues

Fixed Hepa 1-6 tumors were dehydrated using an increasing ethanol concentration and embedded into paraffin to become formalin-fixed paraffin-embedded (FFPE) blocks as previously described.^142^ Following deparaffinization of FFPE samples with xylene and tissue lysis in sodium dodecyl sulfate, total protein was reduced, alkylated, and digested using filter-aided sample preparation ^143^ with sequencing grade modified porcine trypsin (Promega). Tryptic peptides were separated by reverse-phase XSelect CSH C18 2.5 µm resin (Waters) on an in-line 150 × 0.075 mm column using an UltiMate 3000 RSLCnano system (Thermo). Peptides were eluted using a 60 min gradient from 98:2 to 65:35 (buffer A, 0.1% formic acid, 0.5% acetonitrile: buffer B, 0.1% formic acid, 99.9% acetonitrile) ratio. Eluted peptides were ionized by electrospray (2.4 kV) followed by mass spectrometric (MS) analysis on an Orbitrap Exploris 480 mass spectrometer (Thermo). MS data were acquired using a Fourier transform MS (FTMS) analyzer in profile mode at a resolution of 120,000 over a range of 375 to 1500 m/z. Following HCD activation, MS/MS data were acquired using the FTMS analyzer in centroid mode at a resolution of 15,000 and normal mass range with normalized collision energy of 30%. Proteins were identified by database search using MaxQuant (Max Planck Institute) label-free quantification with a parent ion tolerance of 2.5 ppm and a fragment ion tolerance of 20 ppm. Scaffold Q+S (Proteome Software) was used to verify MS/MS-based peptide and protein identifications. Protein identifications were accepted if they could be established with less than 1.0% false discovery and contained at least two identified peptides. Protein probabilities were assigned by the Protein Prophet algorithm.^144^

### RNA-seq data analysis of murine tumors

Hepa 1-6 FFPE scrolls were processed for DNA and RNA extraction using a Quick-DNA/RNA FFPE Miniprep Kit with on-column DNase digestion for the RNA preps (Cat. # R1009; Zymo Research). RNA was assessed for mass concentration using the Qubit RNA Broad Range Assay Kit (Cat. # Q10211; Invitrogen) with a Qubit 4 fluorometer (Cat. # Q33238; Invitrogen). RNA quality was assessed with a Standard Sensitivity RNA Analysis Kit (Cat. # DNF-471-0500; Agilent) on a Fragment Analyzer System (Cat. # M5310AA; Agilent). Sequencing libraries were prepared using TruSeq Stranded Total RNA Library Prep Gold (Cat. # 20020599; Illumina). RNA DV200 scores were used to determine fragmentation times. Libraries were assessed for mass concentration using a Qubit 1X dsDNA HS Assay Kit (Cat. # Q33231; Invitrogen) with a Qubit 4 fluorometer (Cat. # Q33238; Invitrogen). Library fragment size was assessed with a High Sensitivity NGS Fragment Analysis Kit (Cat. # DNF-474-0500; Agilent) on a Fragment Analyzer System (Cat. # M5310AA; Agilent). Libraries were functionally validated with a KAPA Universal Library Quantification Kit (Cat. # 07960140001; Roche). Sequencing was performed to generate paired-end reads (2 × 100 bp) with a 200-cycle S1 flow cell on a NovaSeq 6000 sequencing system (Illumina).

### Bioinformatics analysis

We examined the mRNA and protein expression profiles of Hepa 1-6 tumors treated with PBS or MMR. Three replicates were used to analyze each of the untreated (PBS) and treated (MMR) groups. The tumor samples were sequenced on an NGS platform. The files containing the sequencing reads (FASTQ) were then tested for quality control (QC) using MultiQC.^145^ The Cutadapt tool trims the Illumina adapter and low-quality bases at the end. After the quality control, the reads were aligned to a mouse reference genome (mm10/GRCm38) with the HISAT2 aligner^146^, followed by counting reads mapped to RefSeq genes with feature counts. We generated the count matrix from the sequence reads using HTSeq-count.^147^ Genes with low counts across the samples affect the false discovery rate, thus reducing the power to detect differentially expressed genes; thus, before identifying differentially expressed genes, we filtered out genes with low expression utilizing a module in the limma-voom tool^81^. Then, we normalized the counts by using TMM normalization^148^, a weighted trimmed mean of the log expression proportions used to scale the counts of the samples. Finally, we fitted a linear model in limma to determine differentially expressed genes and expressed data as mean ± standard error of the mean. All *p* values were corrected for multiple comparisons using Benjamini-Hochberg FDR adjustment. After identifying differentially expressed genes, enriched pathways were performed using the Ingenuity Pathway Analyses tool to gain biological insights. The statistical difference between groups was assessed using the nonparametric Mann-Whitney U test R module.

### Integration of transcriptomics and proteomics

The limma-normalized transcript expression levels and the normalized protein intensities were integrated using two independent methods. Firstly, the mixOmics package (Omics Data Integration Project R package, version 6.1.1) was implemented to generate Fig. 6 and Supplementary Fig. 7 as previously described.^149^ Secondly, the MOGSA package was used to generate Supplementary Figs. 8, 9, and 10.^102^

### Murine immunoglobulin isotyping following treatment with MMR

Serum samples were obtained from infected tumor-bearing mice from whole blood collected in BD Microtainer tubes. Mouse anti-measles IgG (Cat. # 530-130-MMG; Alpha Diagnostic), mouse anti-mumps IgG (Cat. #520-130-MMG; Alpha Diagnostic), and mouse anti-rubella IgG (Cat. # 510-120-MRG; Alpha Diagnostic) were used to determine the levels of anti-measles, anti-mumps, and anti-rubella antibodies in the serum samples by ELISA according to the manufacturer’s instructions.

### Blood chemistry and cytokines

Blood chemistry analysis was performed with an Abaxis Piccolo Xpress chemistry analyzer (Abaxis) to assess liver toxicity (i.e., aspartate transaminase, alkaline phosphatase, albumin), nephrotoxicity (i.e., creatinine, blood urea nitrogen), and serum electrolytes. Murine type I interferon-beta assay was performed using Mouse IFN beta SimpleStep ELISA® Kit (Cat. # ab252363; Abcam).

### Quantitative reverse transcriptase polymerase chain reaction

Quantitative reverse transcriptase polymerase chain reaction (qPCR) primers specific to measles nucleocapsid gene, mumps matrix gene, and rubella (envelope glycoprotein E1) were designed and synthesized using the PrimeTime(tm) qPCR program (Integrated DNA Technologies [IDT], USA). First, a reverse transcription-polymerase chain reaction was performed to generate cDNA using the High Capacity cDNA Reverse Transcription Kit (Cat. # 4374966; Applied Biosystems). Followed by amplification of the cDNA to detect the presence of measles, mumps and rubella using the GeneAmp® Fast PCR Master Mix (2X) (Cat. # 4359187; Applied Biosystems). Cycle conditions were 10 minutes at 95 °C followed by 45 cycles of 45 s at 93 °C, 1 minute at 72 °C and 5 minute at at 72 °C. Finally, qPCR reactions were performed on an Applied Biosystems StepOnePlus(tm) Real-Time PCR System (Applied Biosystems) using the PowerUp™ SYBR ™ Green Master Mix (Cat. # A25742; Applied Biosystems). Cycle conditions were 10 minutes at 94 °C followed by 40 cycles of 10 s at 94 °C and 1 minute at 60 °C. Cycle threshold extraction was performed using the iCycler IQ software (version 3, Biorad). The following primers were used: forward primer (measles)-5’CCT CAA TTA CCA CTC GAT CCA G 3’, reverse primer (measles)-5’ TTA GTG CCC CTG TTA GTT TGG 3’; forward (mumps) 5’ TCA AGC CAG AAC AAG CCT AG 3’, reverse (mumps)-5’ TTG ATA ACA GGT CCA GGT GC 3’ and forward (rubella) 5’ TTG AAC CTG CCT TCG GAC 3’, reverse (rubella)-5’ CCT GGT CTC TGT ATG GAA CTT G 3’.

### Statistical analysis

All values were expressed as the mean ± standard error of mean, and the results were analyzed by one-way analysis of variance followed by the Tukey test or Benjamini-Hochberg FDR adjustment for multiple comparisons and *t* test to compare group means. Kaplan-Meier method for survival, using statistical software in GraphPad Prism, version 8 (GraphPad Software). A *p* value less than 0.05 was considered statistically significant.

### Data deposition

The the proteomics data are available via the ProteomeXchange PXD031295.

## ABBREVIATIONS

ALP: alkaline phosphatase
ALT: alanine aminotransferase
Anti-Measles: anti-measles fusion and hemagglutinin antibody
Anti-Mumps: anti-mumps fusion and hemagglutinin antibody
Anti-Rubella: anti-rubella envelope glycoprotein E
AST: aspartate aminotransferase
ATCC: American Tissue Culture Collection
CTLs: cytotoxic T lymphocytes
CRC: Colorectal cancer
DMEM: Dulbecco modified eagle medium
DPBS: dulbecco phosphate-buffered saline
ELISA: enzyme-linked immunosorbent assay
G-MDSCs: granulocytic myeloid-derived suppressor cells
GzmB: granzyme B
h: human
HCC: hepatocellular carcinoma
HGB: hemoglobin
LAV: live attenuated viral vaccine
MOI: multiplicity of infection
PBS: phosphate-buffered saline
PD-1: programmed cell death protein 1
PE: phycoerythrin
RNA: ribonucleic acid
TAMs: tumor-associated macrophages
IT: intratumoral

## Acknowledgments

We thank the personnel of the DNA Damage and Toxicology, Proteomics, Genomics, and Bioinformatic Cores at the University of Arkansas for Medical Sciences (UAMS) for their assistance during these studies.

## Author contributions

BMN, AB and MJC contributed to study concept and design, data acquisition, data analysis, data interpretation, and manuscript drafting. YZ, MG, MT, JCC, CD, OB, MJB, SRP, CSS and MJC contributed to data acquisition, data analysis, data interpretation, drafting and critical revision of the manuscript. All authors approved the final, submitted version of the manuscript.

## SUPPLEMENTARY FIGURES AND TABLES

**Supplementary Figure 1**. *Overview of treatment schedule*. Subcutaneous syngeneic tumor-bearing (Hepa 1-6, MC38) mice were injected intratumorally with three low doses (1 × 10^2^ TCID_50_) of measles, mumps, and rubella (MMR) live vaccine. Tumor and survival for recorded for the two models (Hepa 1-6: 28 days; MC38: 34 days).

**Supplementary Figure 2**. *Analysis of markers of hepatotoxicity and nephrotoxicity following MMR-based intratumoral immunotherapy*. Changes in the markers of liver toxicity (i.e., serum amylase), nephrotoxicity (i.e., creatinine), and blood electrolytes (i.e., calcium) between groups administered measles, mumps, and rubella (MMR) live vaccine or phosphate-buffered saline (PBS; controls) were plotted in graphs.

**Supplementary Figure 3**. *Body weight of tumor-bearing mice treated with MMR vs. PBS*. Graphs showing the effect of treatment with multiple low doses of measles, mumps, and rubella (MMR) live vaccine or phosphate-buffered saline (PBS; controls) on body weights of individual mice with Hepa 1-6 **(a)**, or MC38 **(b)**.

**Supplementary Figure 4**. *Quantification of virus-specific Immunoglobulin in serum*. At the end of the study, Hepa 1-6 (28 days) and MC38 (34 days) mice administered either administered measles, mumps, and rubella (MMR) live vaccine or phosphate-buffered saline (PBS; controls) were sacrificed. Blood was processed to serum to measure the levels of specific immunoglobulin (IgG) to **measles (a)** (negative < 8 U/mL; positive > 10 U/mL), mumps **measles (b)** (negative < 17 IU/mL; positive > 20 IU/mL), and rubella **measles (c)** (negative < 10 IU/mL; equivocal 10-15 IU/mL; positive > 15 IU/mL).

**Supplementary Figure 5**. *Plaque assay for MMR vaccine and amplification of virus-specific genes in tumor tissues*. **a**) A low dose of MMR (1x 10^2^ TCID_50_) was used to infect a monolayer of Vero cells (2.5 × 10^5^); four days later, they were stained with crystal violet to reveal viral plaques. **b)** Specific primers to measles nucleoprotein (N), mumps matrix (M), and rubella envelope protein (E1) genes were used to detect and quantify viral replication in Hepa 1-6 and MC38 tumors after intratumoral immunotherapy based on MMR.

**Supplementary Figure 6**. *Immunophenotyping of Tumor-Infiltrating Immune Cells Following Intratumoral Administration of MMR*. Immune profiling of syngeneic mice administered with PBS or MMR, showing percentages of tumor-infiltrating immune cells (CD3+, total CD8+, CD8+ Ki67, CD8+ CD44+, CD8 IFNg+, CD8+ PD-1+, CD4+ Ki67+, CD4+ CD44+, total CD4+, CD4+ PD-1+, CD4+ IFNg+, total macrophages, NKT, CD11+ in CD45+, M2 macrophages, NK cells and CD11-in CD45+ cells). Bartlett test, ANOVA and a two-sample t-test were used to compare group means.

**Supplementary Figure 7**. *The top 30 most enriched Kyoto Encyclopedia of Genes and Genomes pathways*. KEGG analysis of groups treated with multiple low doses of measles, mumps, and rubella (MMR) live vaccine or phosphate-buffered saline (PBS; controls) showing the top 30 genes involved in MMR-mediated activation of immune response pathways.

**Supplementary Figure 8**. *Genes contributing to effects seen on protein and mRNA results Pathways*. Maps of mRNAs (left) and proteins (right) differentially expressed used to generate the integrated datasets.

**Supplementary Figure 9**. *Heatmap with a filtered by biologically relevant top KEGG pathways that are differential between PBS and MMR*. Consistently enriched biological pathways in all MMR or PBS groups samples were filtered and used to generate a heatmap.

**Supplementary 10**: *Contribution of individual genes to the KEGG pathway’s effect on the phenotype (PBS vs. MMR)*. The gene influential score (GIS) represents the influence of individual genes on the gene-set score (the gene-set score was used to create the heatmap in Supplementary Fig. 8. This metric indicates which dataset (or pathway) contributed most to the biological process (phagocytosis and cellular death).

**Supplementary Figure 11**. *Top MSigDB pathways in the integrated datasets*. Heatmap showing molecular signatures database (MSigDB) enrichment in MMR versus PBS samples.

