## Supplementary Figures for "Repurposing Live Attenuated Trivalent MMR Vaccine as Cost-effective Cancer Immunotherapy"

### Supplementary data

**Supplementary Figure 1**. Treatment schedule used in this study.

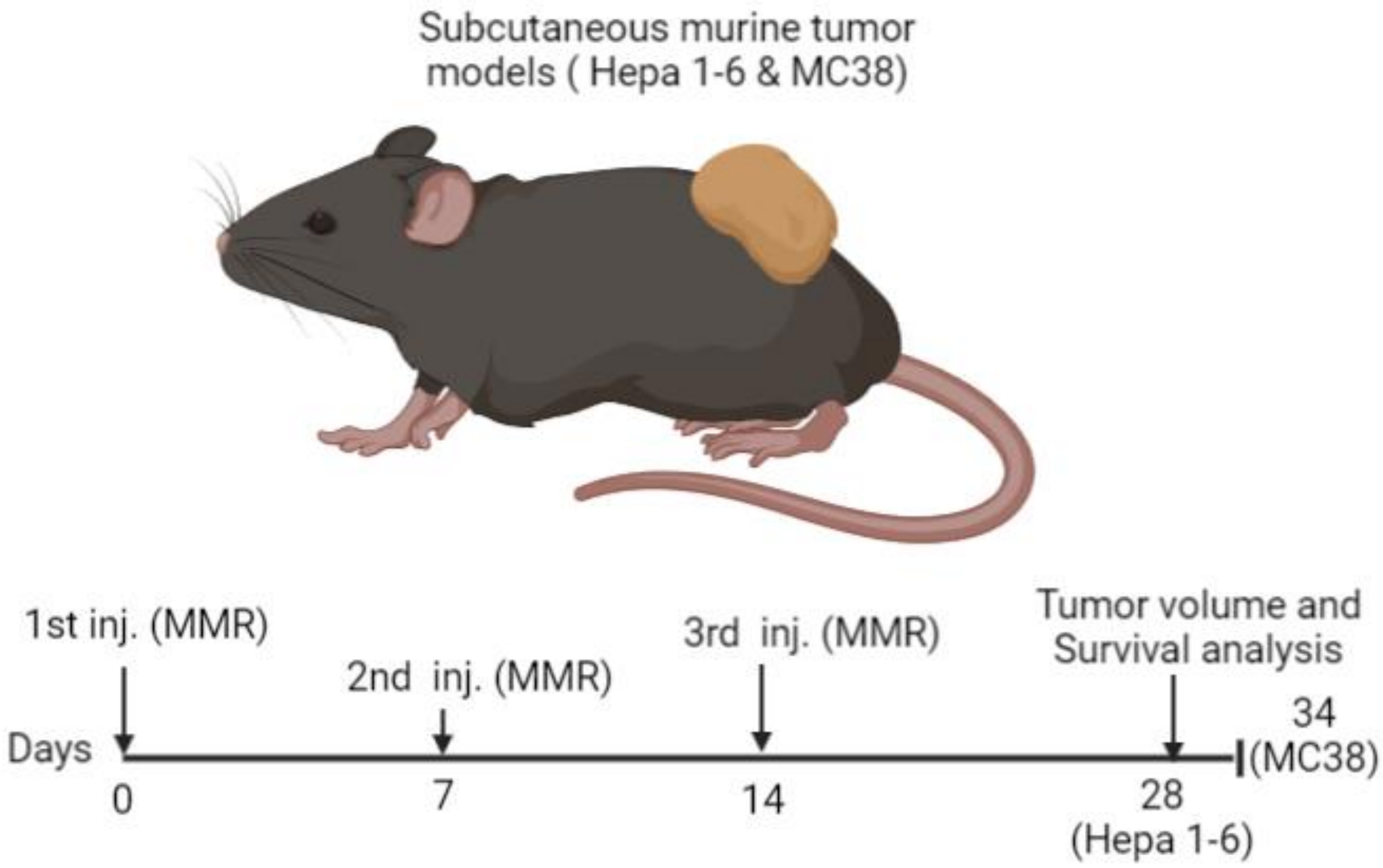

**Supplementary Figure 2**. Analysis of markers of hepatotoxicity and nephrotoxicity following MMR-based intratumoral immunotherapy

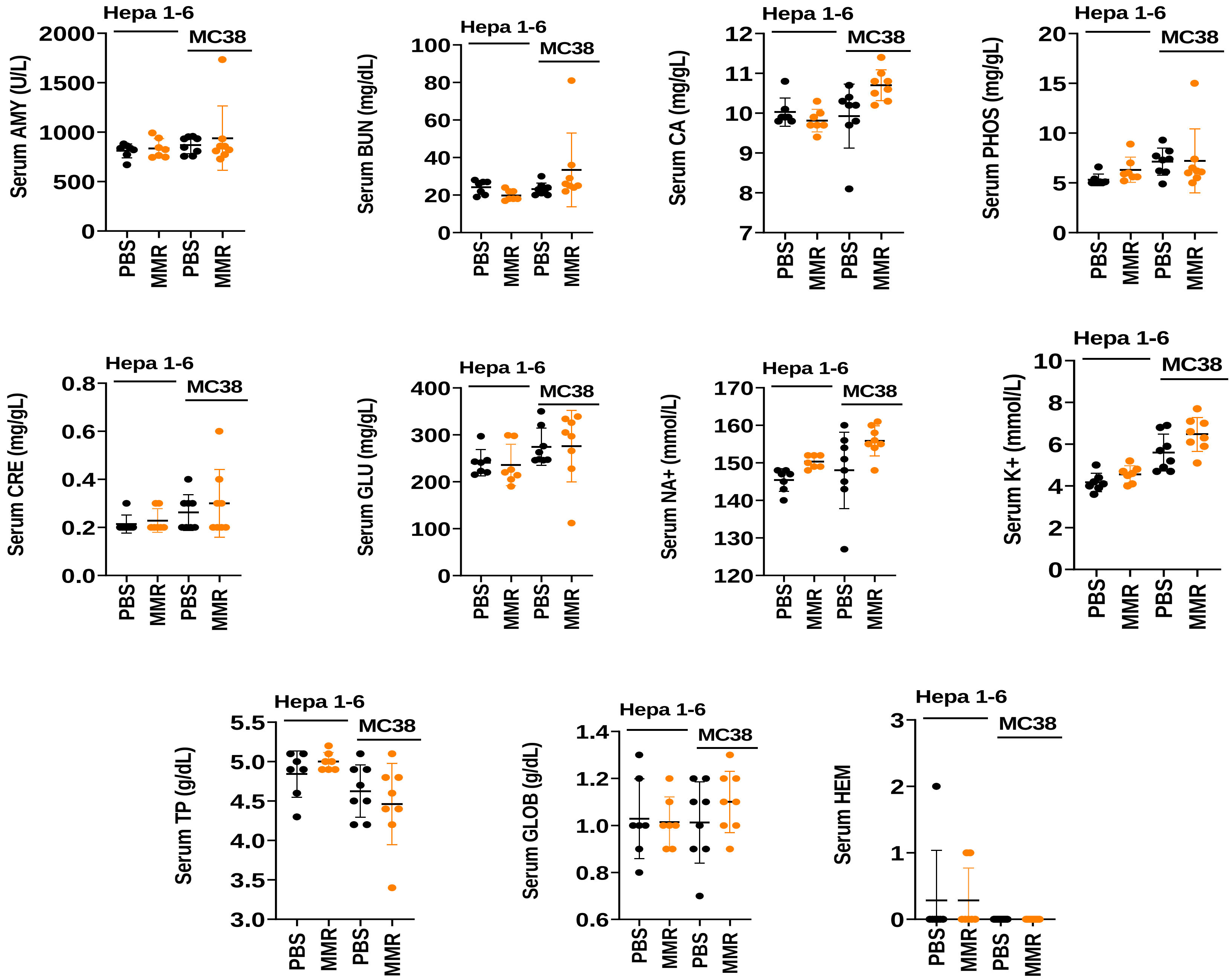

**Supplementary Figure 3**. Body weight of tumor-bearing mice treated with MMR vs. PBS

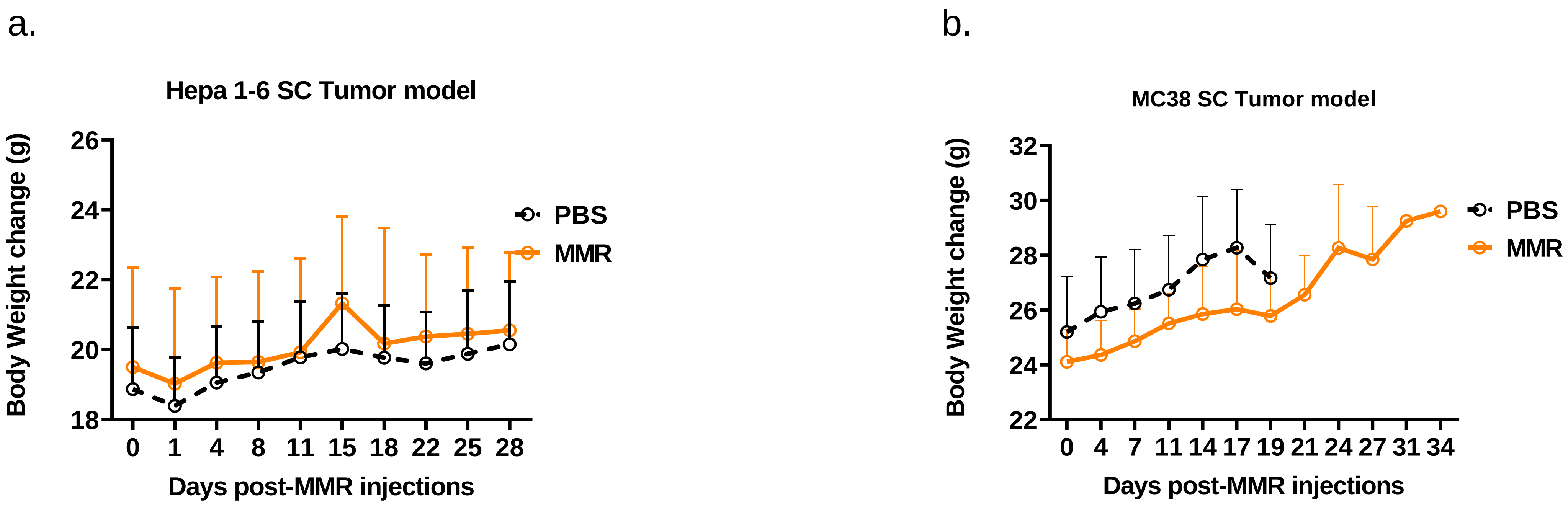

**Supplementary Figure 4**. Quantification of virus-specific Immunoglobulin G (IgG) in serum

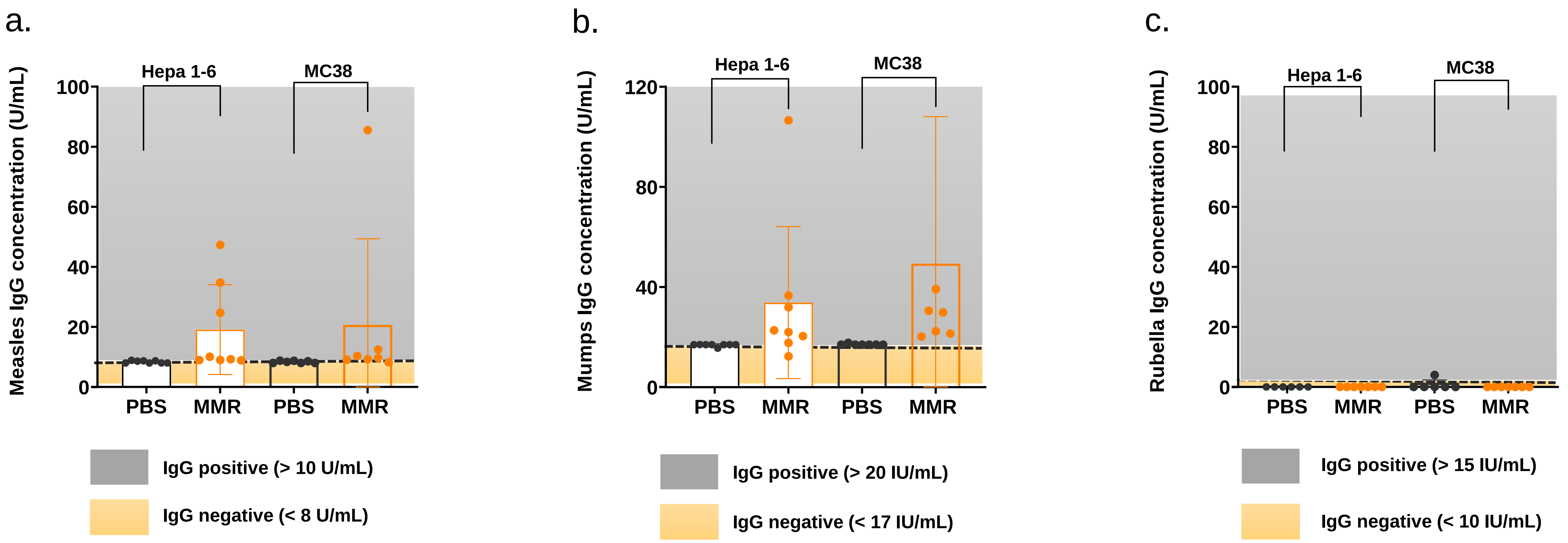

**Supplementary Figure 5**. Amplification of virus-specific genes in tumor tissues

a.

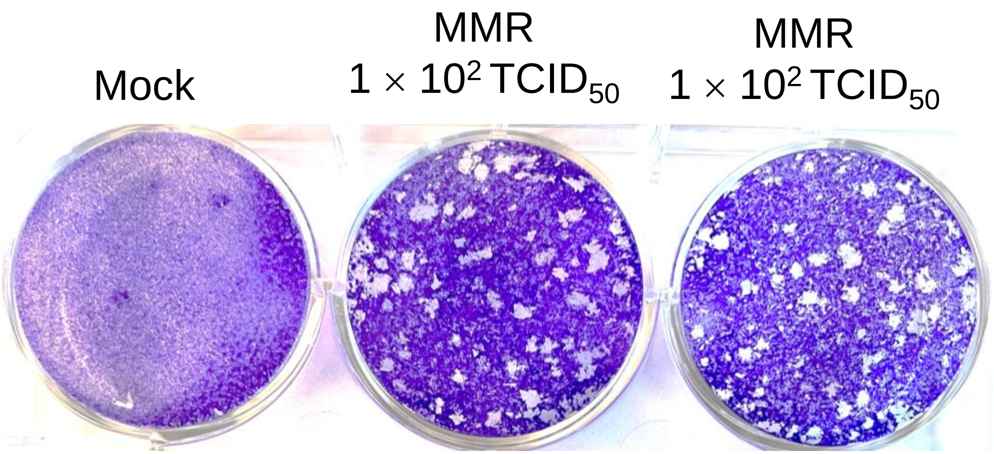


b.

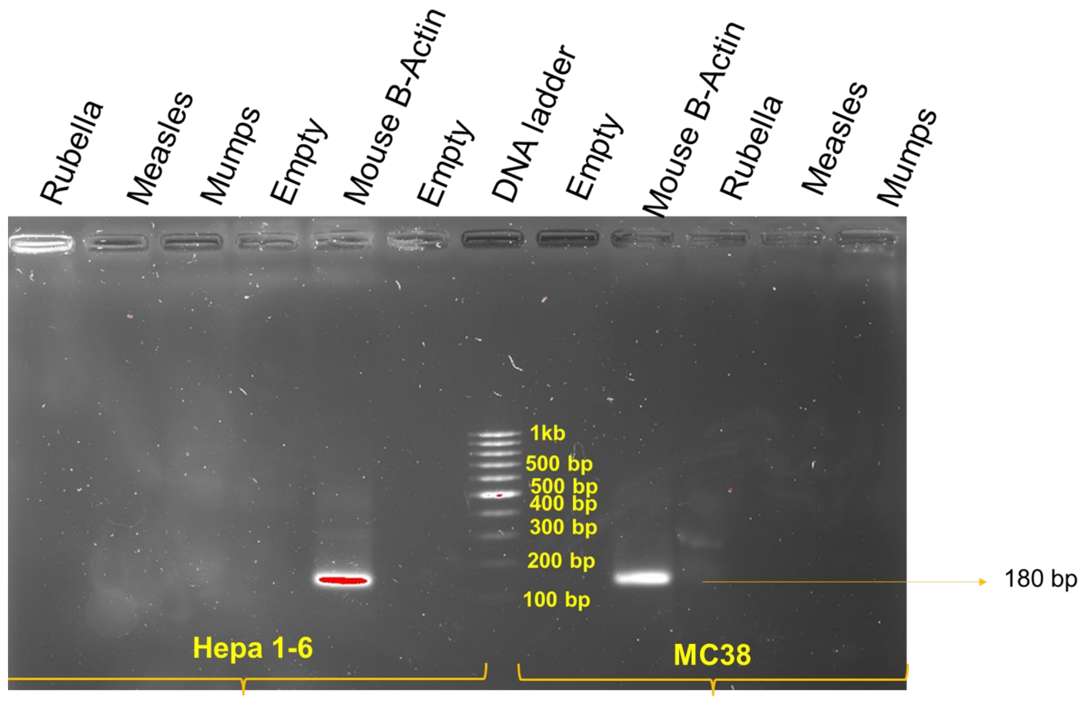

**Supplementary Figure 6.** Immunophenotyping of Tumor Infiltrating Immune Cells Following Intratumoral Administration of MMR

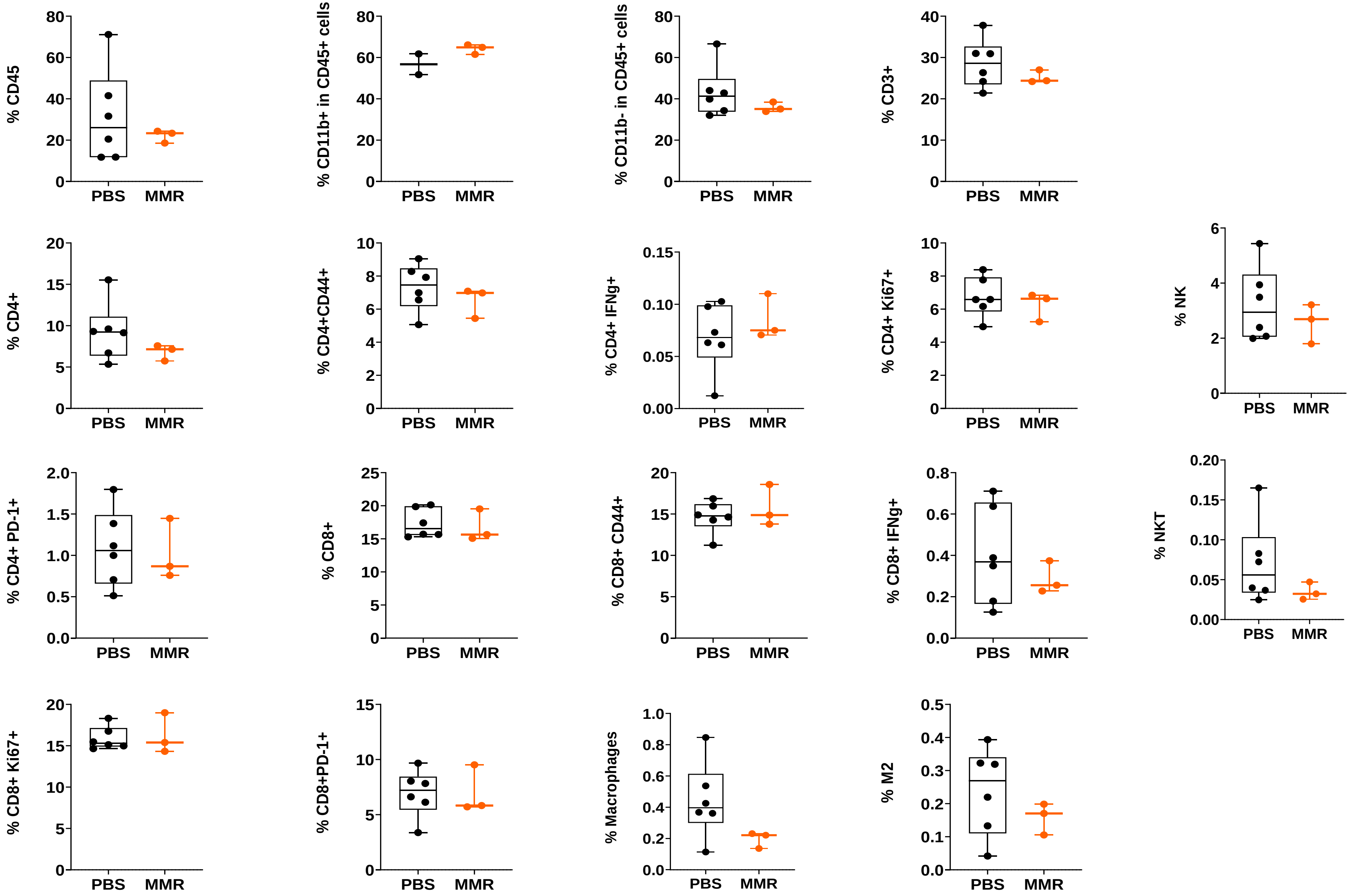

**Supplementary Figure 7.** The top 30 most enriched Kyoto Encyclopedia of Genes and Genomes pathways

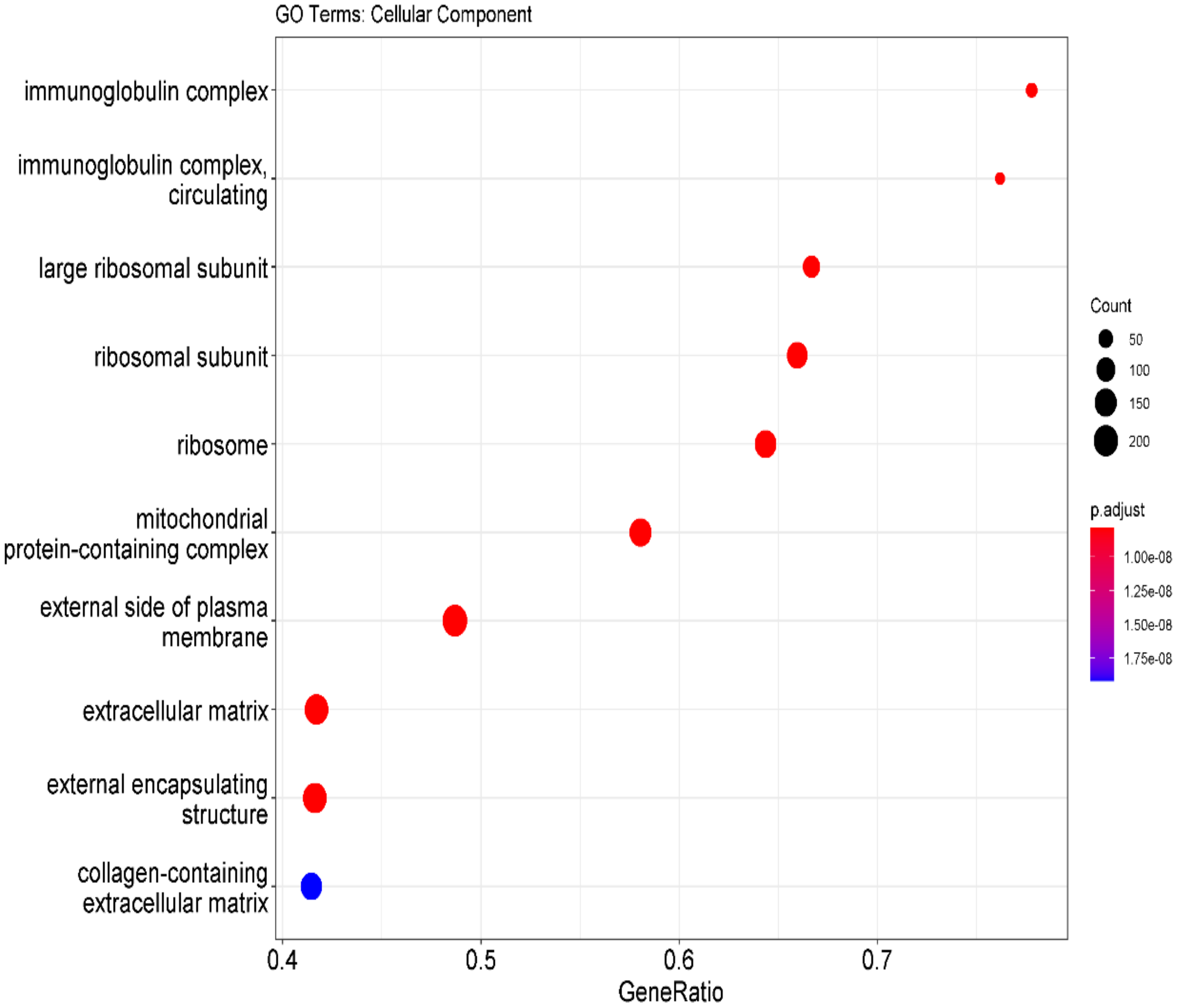

**Supplementary Figure 8.** Genes contributing to effects seen on protein and mRNA results Pathways

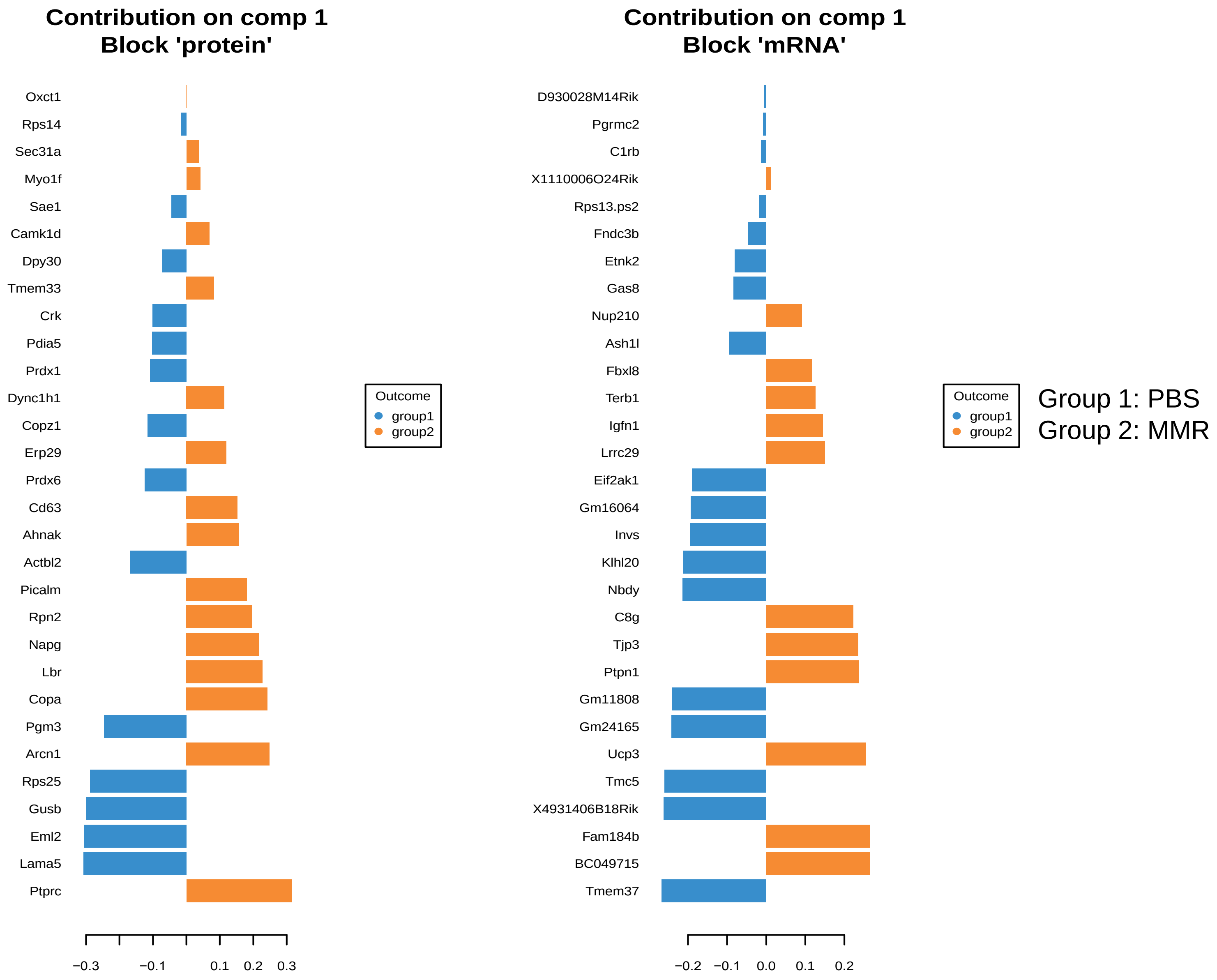

**Supplementary 9**: Heatmap with a filtered top KEGG pathways that are differential between PBS and MMR

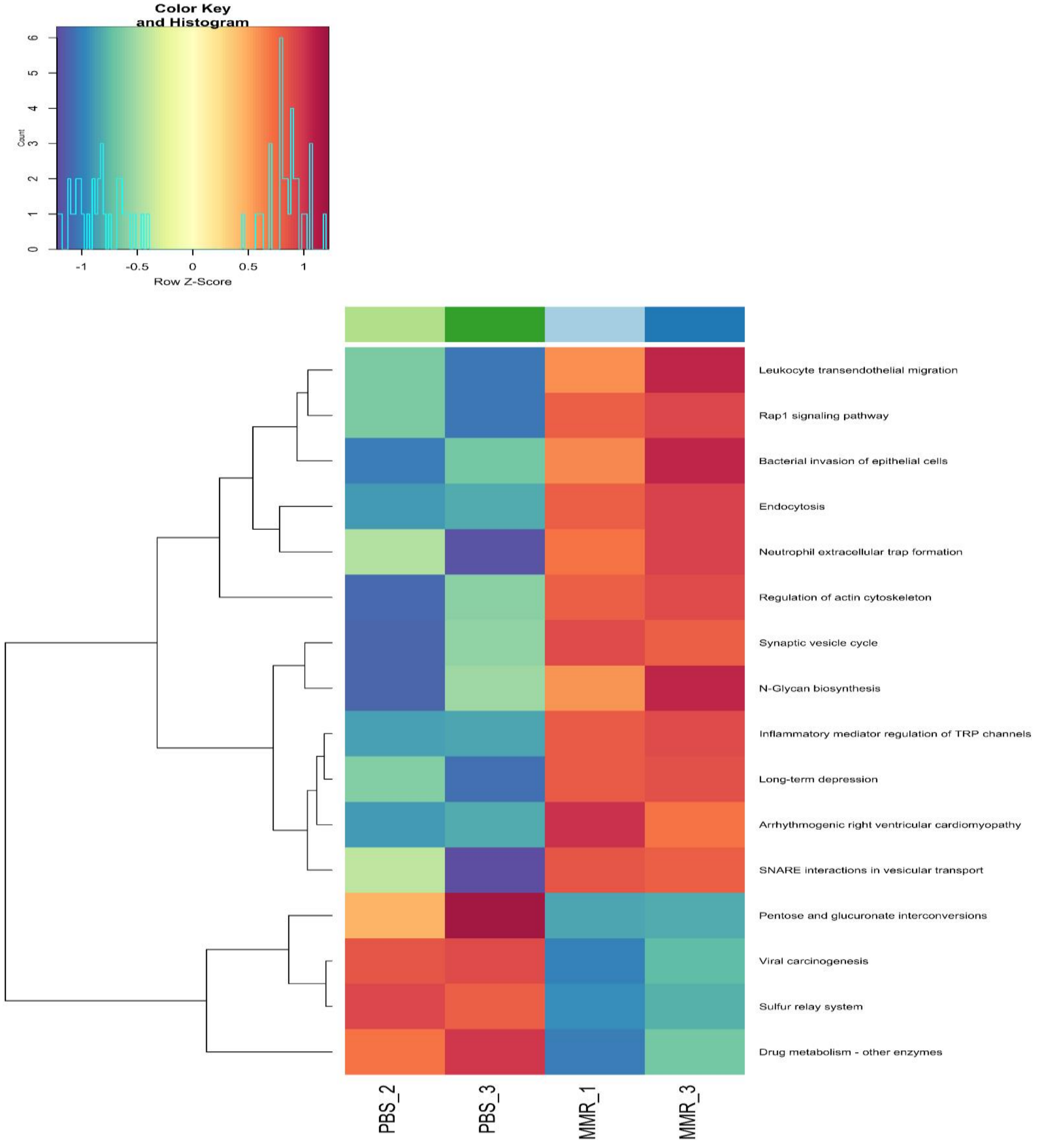

**Supplementary 10**: Contribution of individual genes to the KEGG pathway's effect on the phenotype (PBS vs MMR)

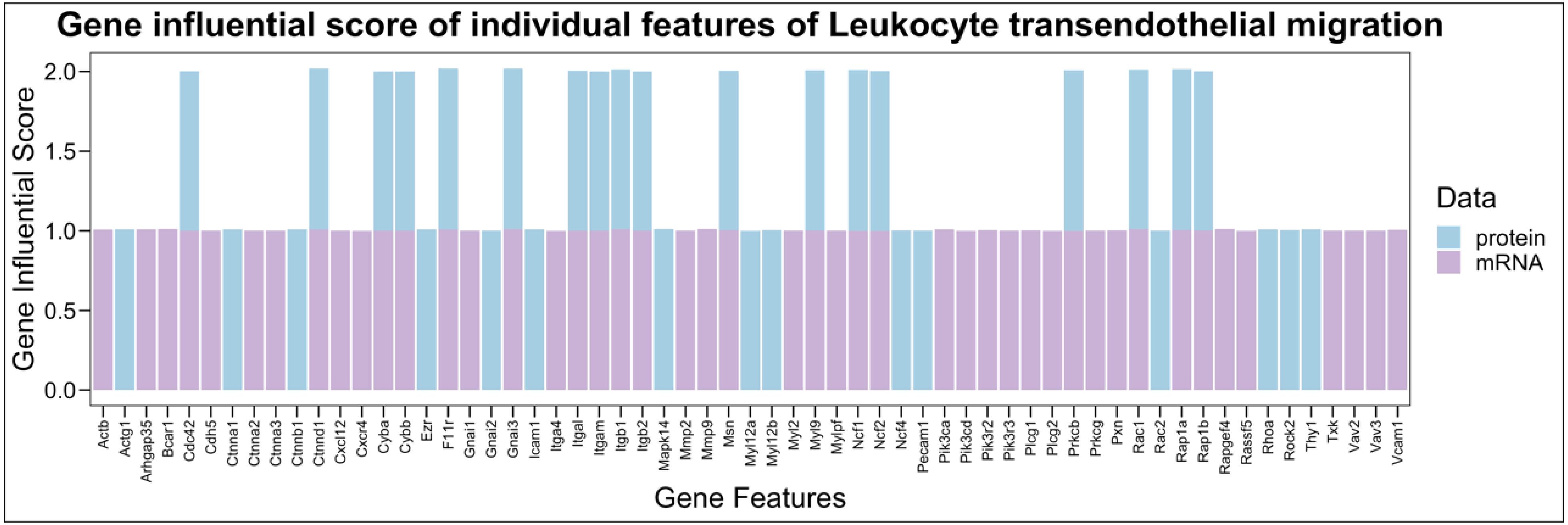


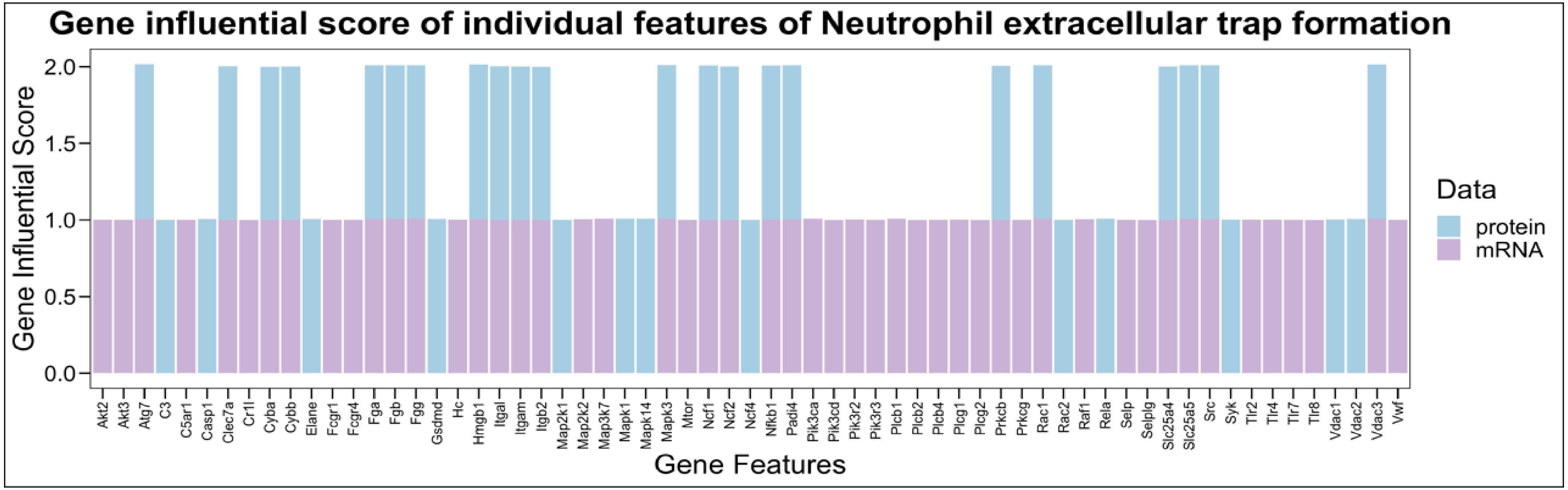

**Supplementary Figure 11.** Top MSigDB pathways in the integrated datasets

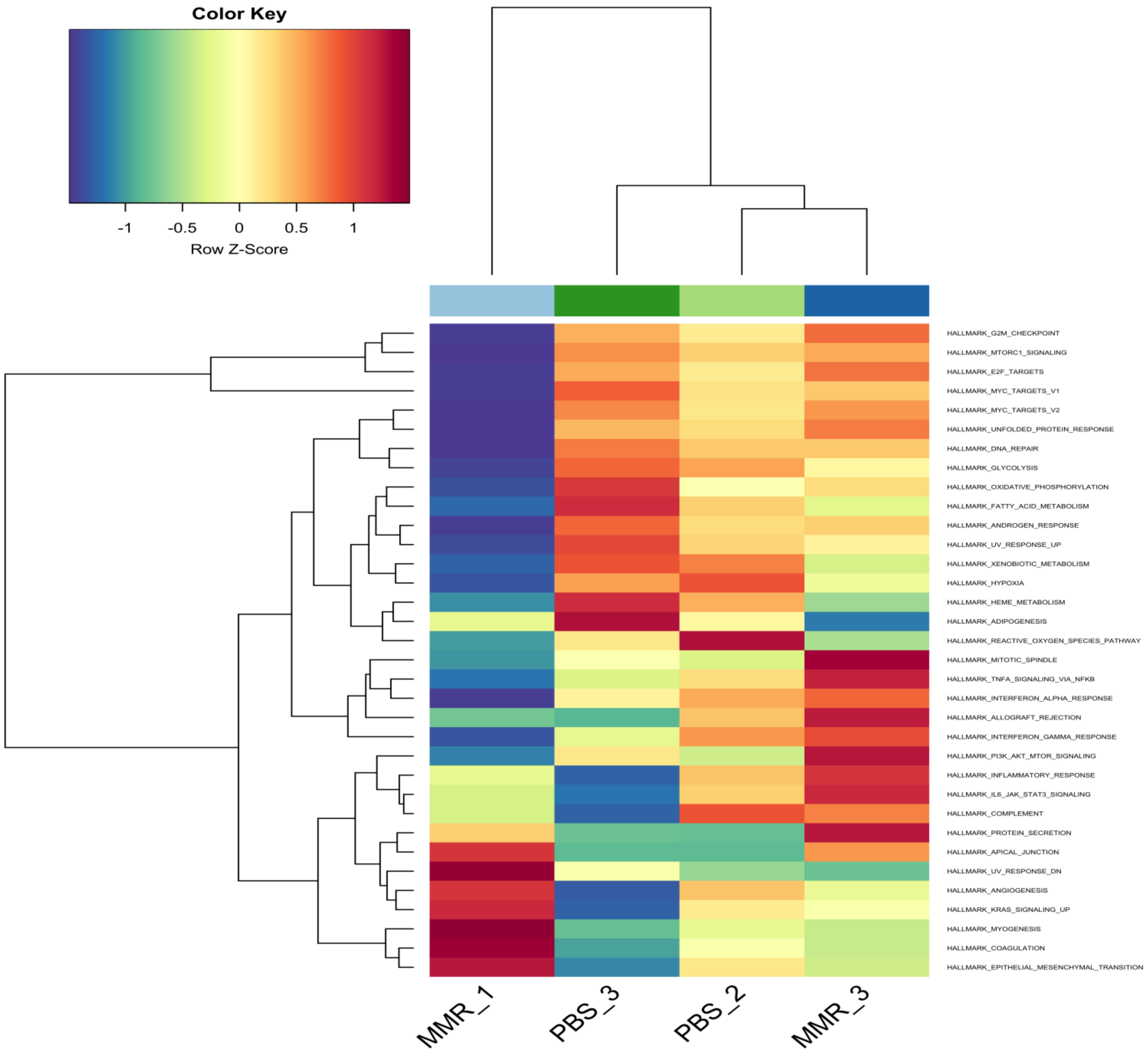
